# Development of a basal media for carbon utilization assay and analysis of the starch utilization locus in *Prevotella melaninogenica*

**DOI:** 10.1101/2025.07.04.663206

**Authors:** Claire E. Albright, Kelyah Spurgeon, Souzane Ntamubano, Ariangela J. Kozik

## Abstract

*Prevotella melaninogenica* is a highly abundant member of the human respiratory microbiota, occupying a range of distinct niches from the oral cavity to the lower airways. Colonization within these nutrient diverse environments suggest a sophisticated metabolic system equipped to deal with large fluctuations in nutrient availability. However, the core physiology and metabolic profile of this bacteria remain largely uncharacterized. In this study, we developed a basal media conducive to assaying carbon utilization of *P. melaninogenica* and demonstrate the utilization of dietary carbohydrates with varied glycosidic bonds, including the starch mimic cyclodextrin. Furthermore, we identified the putative starch utilization system, highlighting the presence of three novel components which are absent from starch metabolism in the closely related *Bacteroides* genus. Together, these findings provide the first characterization of the nutritional profile and central metabolic capacity of *P. melaninogenica*, offering crucial insight into how this bacterium adapts to and thrives within ecologically distinct niches.

**IMPORTANCE:** This work significantly advances our understanding of the metabolic capabilities of Prevotella melaninogenica, a common yet largely unexplored member of the respiratory microbiota. By mapping key aspects of its predicted starch utilization locus, we lay the foundation for future studies on microbial adaptation and persistence in the respiratory tract.

## INTRODUCTION

*Prevotella melaninogenica* is saccharolytic, Gram negative bacterium, and appears as non-motile short rods or coccobacilli (1). *P. melaninogenica* was first isolated by Oliver and Wherry in 1921 from various human body sites and was named for the black pigmentation of its colonies when grown on blood agar. The advent of cultivation-free microbial profiling showed that *P. melaninogenica* is highly prevalent and abundant in the human respiratory tract, spanning diverse habitats from the oral cavity down to the lungs. In the mouth, *P. melaninogenica* is estimated to make up roughly 4-8% of the oral microbiome (2), and in the lower respiratory tract (which is seeded by the mouth), it has been found by our group to make up a staggering 5-35% (unpublished data). These findings strongly suggest that *P. melaninogenica* plays a large role in the microbial ecology throughout the human respiratory tract.

*P. melaninogenica* has been reported to be associated with both the healthy respiratory system (3) and various respiratory diseases such as long-term COVID, Gastrointestinal Reflux Disease (4,5), COPD (6) and Cystic Fibrosis (7). However, *P. melaninogenica* (and other *Prevotella spp.)* has received significantly less attention in comparison to other members of the human microbiome. This is likely due to the challenging nature of culturing this oxygen-sensitive bacteria(8). This methodological hurtle has in turn impeded advances in the direct study of its impact on human respiratory health and disease, resulting in a stall at *association* level findings.

Nevertheless, the successful colonization of *P. melaninogenica* within strikingly varied environments demonstrates both its relevance to microbiome community dynamics and the human host. Therefore, the adaptive mechanisms of *P. melaninogenica* to the host is an important first step in elucidating the impact of this bacteria on microbiome ecology and human health.

In this study we show that *P. melaninogenica* can uptake and metabolize dietary carbohydrates with varied glycosidic bonds, including the starch mimic cyclodextrin. Furthermore, we identified the putative starch utilization system, highlighting the presence of three novel components which are absent from starch metabolism in the *Bacteroides* genus. Together, these findings begin to elucidate the nutritional profile and mechanism of metabolism in *P. melaningenica*, offering critical insights into how this bacterium can successfully colonize niches with vastly different nutritional availability and microbial communities.

## RESULTS

### Development of a Basal Media for metabolic assay in *P. melaninogenica*

Our first objective was to develop a media which yields a significant positive change in *P. melaninogenica* growth (hereafter referred to as +Δ) when supplemented with carbon. This work is crucial as the existing medias for carbon utilization assay are either not conducive to *P. melaninogenica* growth or do not produce a significant +Δ with carbon supplementation (Fig 1a). To develop a bacterial basal media (BBM), we utilized low levels of peptides along with the essential growth factors hemin, vitamin K, and menadione (Table 1). We chose to utilize the dietary monosaccharides a-D-glucose and D-fructose as carbon sources for supplementation testing due to their relevance in central metabolism. A control of *P. melaninogenica* in BBM without carbon supplementation was included for each experiment. The resulting control showed the desired low-level growth with a maximum optical density (OD) of 0.26. In contrast, when supplemented with a-D-glucose, an average max OD of 0.71 was measured. This produces a +Δ of 0.45 in the max OD, indicating a significant increase in growth with glucose. Similarly, when supplemented with D-fructose, a max OD of 0.61 and subsequent +Δ of 0.35 in the max OD was measured (Fig 1b). These findings demonstrate that our newly developed BBM media is conducive to the quantification of carbon source utilization in P. *melaninongenica*.

**Figure 1.**
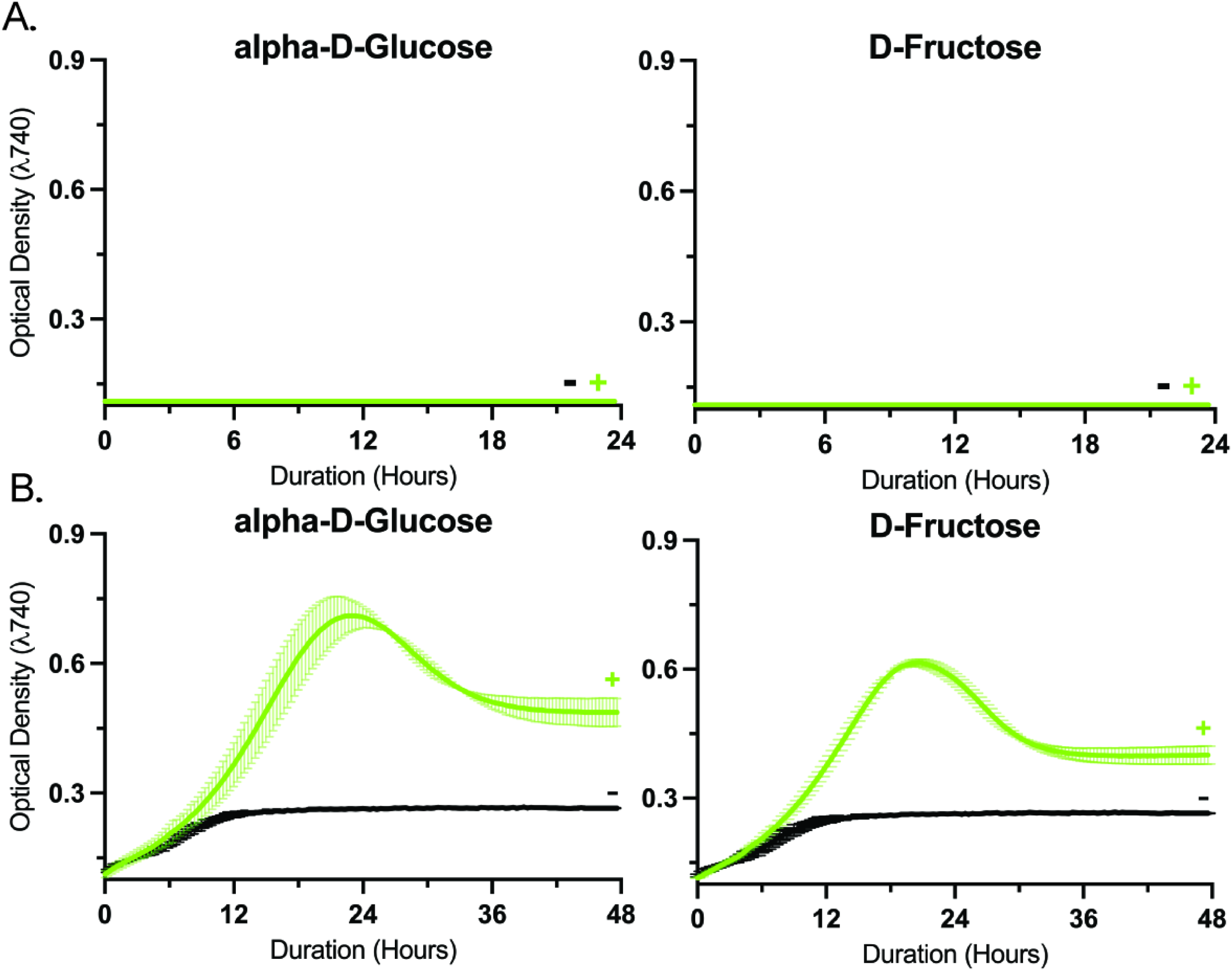
Creation of a basal media for *P. melaninogenica*. Growth curves with supplementation are represented by green curves and “+”. Negative control cultures without additional supplementation are labeled with “-”. A. *P melaninogenica* +/- Glucose and Fructose suppllementation in a commer­ cial carbon assay media. B. *P melaninogenica* +/- Glucose and Fructose supple­ mentation in our newly developed Bacterial Basal Media.

**Table 1.**
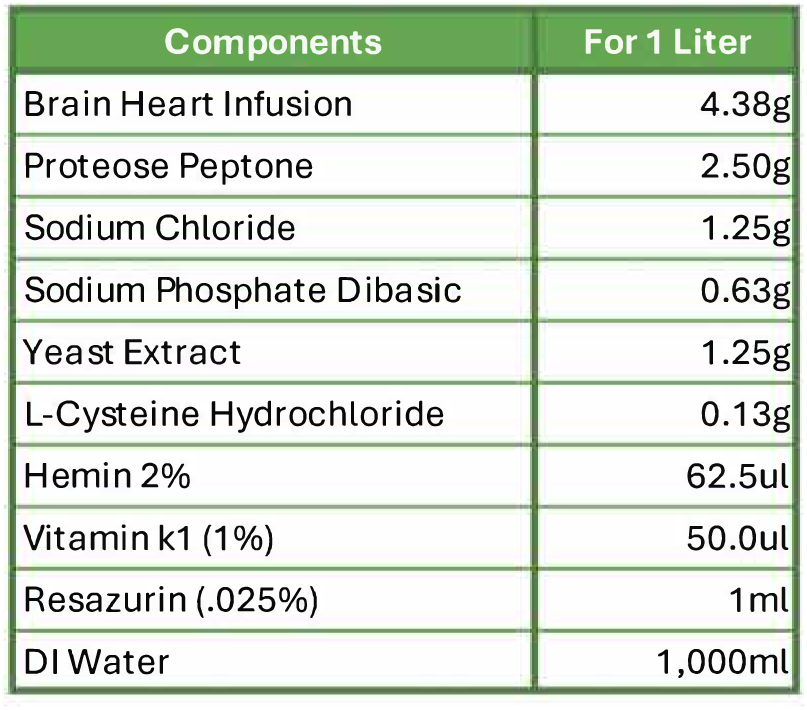
Bacterial Basal Media Recipe. Our developed recipe of Bacterial Basal Media (BBM). Componenets are prepared according to the methods section.

*P. melaninogenica* can hydrolyze α 1,4, β 1,4 and α 1,2 bonds but cannot hydrolyze α and β 1,6 bonds for glucose utilization.

Our objective was to determine whether *P. melaninogenica* can hydrolyze a variety of glycosidic linkages in glucose-containing di-, tri-, and oligosaccharides for subsequent utilization in central metabolism. As it is likely that *P. melaninogenica* may metabolize free glucose, we focused on carbohydrates composed of glucose monomers. To assess substrate utilization, we quantified bacterial growth in the presence and absence of common dietary carbohydrates featuring diverse glycosidic bonds.

We first tested a panel of starch derivatives which only contain α 1,4 bonds and glucose monomers (Fig 2a). Supplementation with the disaccharide D-maltose, the second smallest unit of starch after glucose, yielded the greatest increase in *P. melaninogenica* growth and utilization rate of all the starch derivatives tested. Compared to the negative control, a +Δ max OD of 0.57 and a replication rate of 6.15 hours was measured. Supplementation with the trisaccharide maltotriose and the cyclic olisaccharides α- and γ-cyclodextrin were all found to yield similar +Δ max ODs of between 0.43 and 0.46. However, while the replication rate with maltotriose was like that of D-maltose at 6.6 hours, supplementation with α- and γ-cyclodextrin both yielded significantly slower replication rates of 14.15 and 13.19 hours respectively. Therefore, *P. melaninogenica* can hydrolyze the α 1,4 glycosidic linkage and utilize these starch derivatives in central metabolism, however the rate of metabolism varies between smaller molecular weight compounds and the larger cyclic compounds.

**Figure 2.**
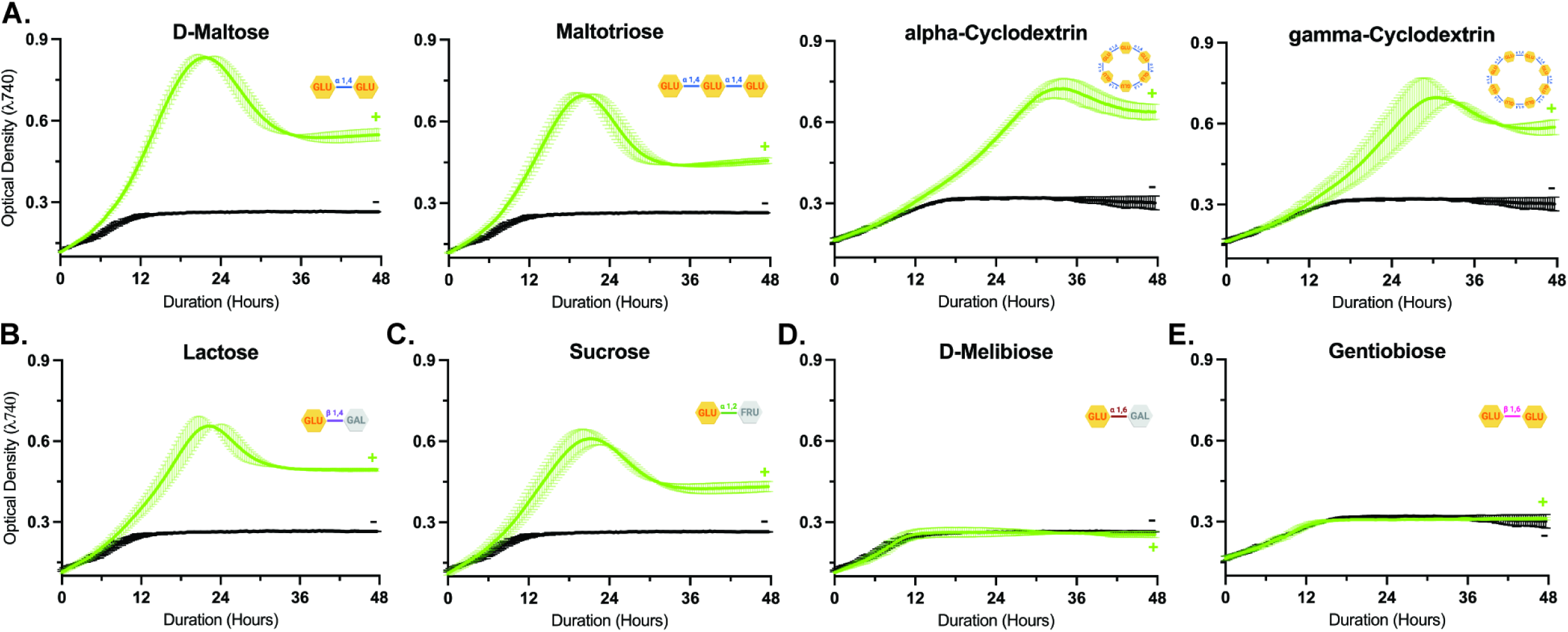
Carbon source utilization profile in *P melaninogenica* indicates enzymatic potential for glycosidic bond hydrolysis. Growth curves of *P melaninogenica* grown in Bacterial Basal Media and supplemented with a panel of carbon sources. Within each graph, cultures grown with carbon supplementation are represented as a green curve notated with a “+”. The negative control growth curves in black, grown in the absence of additional supplementation, are notated with a“-”. The optical density was measured at wavelength 740 for all samples. Measurements were taken *every* ten minutes for 48 hours. A. Sugars containing alpha 1,4 glycosidic bonds. The disaccharide D-Maltose, trisaccharide Maltotriose, and starch mimics alpha-Cyclodextrin and gamma-Cyclodextrin consist only of glucose monomers. B. Lactose, a sugar consisting of glucose and galactose monomers linked by a beta 1,4 bond. C. Sucrose, a sugar consisting of glucose and fructose monomers linked by an alpha 1,2 bond. D. O-Melibiose, a sugar consisting of glucose and galactose linked by an alpha 1,6 bond. E. Gentiobiose, a sugar consiting of two glucose monomers linked by a beta 1,6 bond.

We next tested the β 1,4 bond containing disaccharide lactose, a compound made up of a glucose and galactose monomer (Fig 2b). Lactose demonstrated a +Δ max OD of 0.44 and a replication rate of 8.15 hours. This replication rate is slower than maltose and maltotriose but faster than the cyclodextrins. Furthermore, the lactose metabolizing enzyme β-galactosidase (EC:3.2.1.23) was predicted to be present in the *P. melaninogenica* 25845 genome at the locus tag HMPREF0659_RS11010. These results indicate that *P. melaninogenica* can hydrolyze the β 1,4 bond and use the glucose monomers for increased growth. Interestingly, the α-1,2 bond containing disaccharide sucrose (consisting of a glucose and fructose monomer), had a slightly lower +Δ max OD of 0.34 but a slightly faster replication rate of 7.20 hours (Fig 2c). The sucrose metabolizing enzyme invertase (EC 3.2.1.26) was predicted to be present in the *P. melaninogenica* 25845 genome at the locus tag HMPREF0659_RS00640. Therefore, this suggests that *P. melaninogenica* can hydrolyze the β 1,4 and α-1,2 bonds at slightly varying rates but with no significant difference in utilization.

To test the hydrolysis of α and β 1,6 bonds, *P. melaninogenica* growth was measured +/- D-melibiose or gentiobiose respectively (Fig 2d-e). The resulting data demonstrated no increase in growth when supplemented with D-meliobiose and gentiobiose.

Interestingly, when examining the genome, both alpha-glucosidase (EC 3.2.1.20) and endo-beta-1,4-glucanase (EC 3.2.1.4), the enzymes classically found to hydrolyze these bonds, were predicted to be encoded at HMPREF0659_RS00515 and HMPREF0659_RS02165.

Overall, these findings demonstrate *P. melaninogenica*’s ability to hydrolyze α 1,4 bonds and utilize starch-derived compounds of varying complexity (with the rate of utilization potentially coinciding with compound size). Furthermore, *P. melaninogenica* can hydrolyze β 1,4 and α-1,2 bonds to utilize the glucose within fructose and sucrose respectively. However, *P. melaninogenica* is unable to utilize D-meliobiose and gentiobiose, suggesting its inability to hydrolyze α and β 1,6 bonds.

### The putative glycolytic genes in *P. melaninogenica* depict a complete glycolytic pathway

Our objective was to determine if the genes necessary for the breakdown of glucose to pyruvate were present in *P. melaninogenica*. Due to the lack of a genetically tractable strain of *P. melaninogenica*, we turned to bioinformatics as our method of analysis. We first utilized the KEGG pathway database to identify glycolytic genes predicted to be present and absent in *P. melaninogenica.* We then cross compared these results to the genes present in *P. melaningoenica* ATCC 25845 using the genome annotation software DRAM (which pulls from KEGG, UniRef90, and MEROPS databases) to confirm their presence or absence in our strain. We found that all the genes necessary for the breakdown of glucose to pyruvate were present (Fig 3). Notably, many of the shunts typically present in central glycolysis were absent in *P. melaninogenica* (Sup 2). In conclusion, *P. melaninogenica* is predicted to contain all necessary genes for central glycolysis.

**Figure 3.**
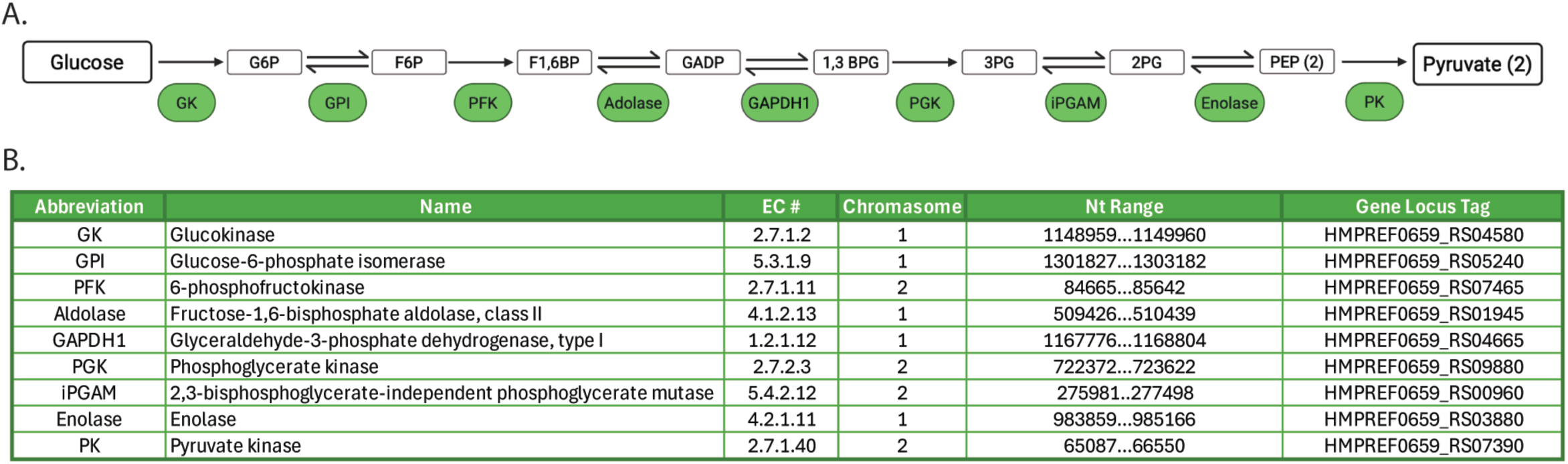
Glycolytic genes predicted to be present in *P melaninogenica* depict a complete glycolytic pathway. A. Model of the central glycolytic pathway in *P melaninogenica,* ommiting shunts and enzymes with redundant function. Enzymes are shown in green. B. A table of the glycolytic enzymes, their EC number, chromosome, nucleotide range and gene locus tag.

### Gene cluster PUL3 in *P. melaninogenica* encodes the putative starch utilization system

To identify the genes facilitating starch (and starch mimic cyclodextrin) uptake and breakdown in *P. melaninogenica*, we first aimed to identify its predicted polysaccharide utilization loci (PUL) (Fig 4). Our approach was to first utilize the PUL database (PULdb) to identify candidate gene clusters. The PULdb predicts the presence of PULs in a genome based on the presence of two conserved PUL genes*: susC* and *susD*. We found 25 predicted PULs in *P. melaninogenica*.

**Figure 4.**
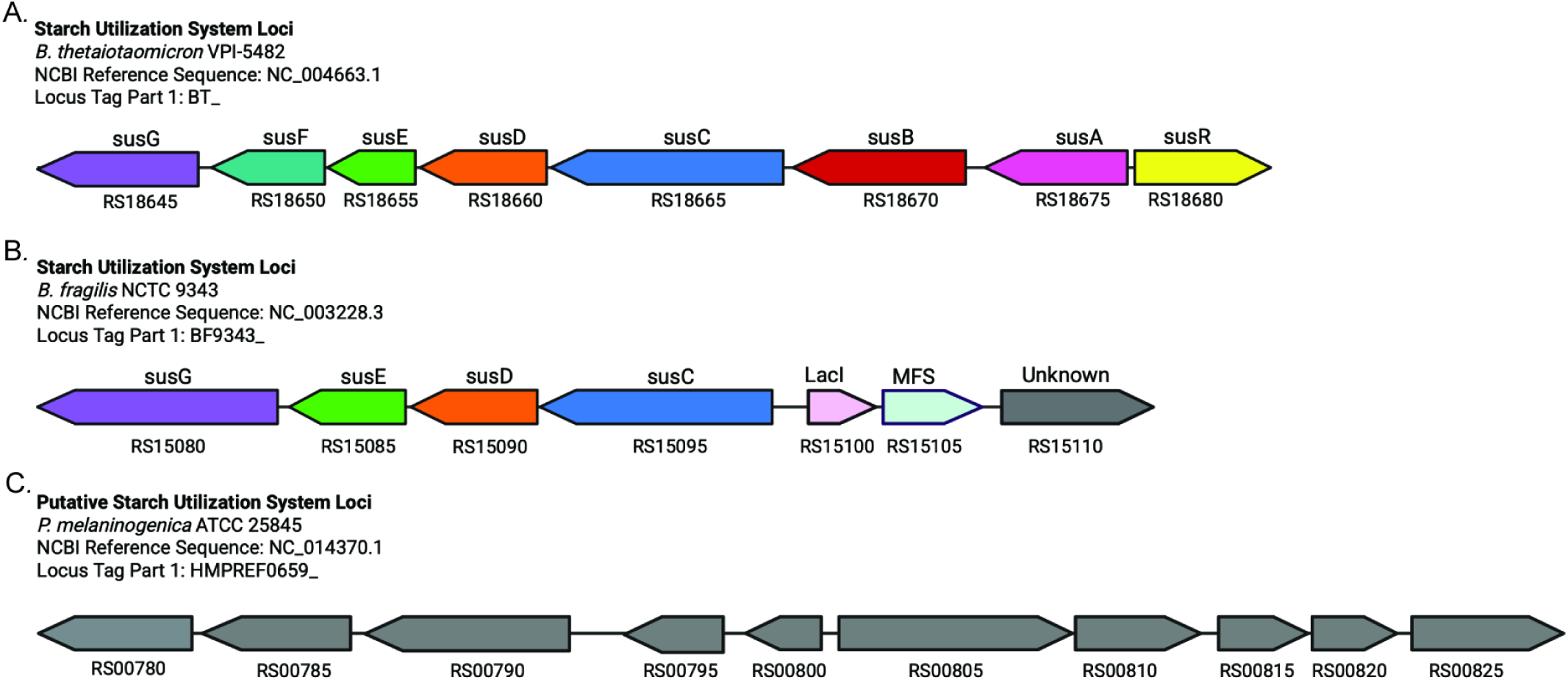
The putative starch utilization system gene cluster in *P melaninogenica*. A. *B. thetaiotaomicron’s* well characterized starch utilization system (sus) loci consisting of an eight gene cluster. B. *B. fragilis’s* seven gene cluster sus loci. C. *P melaninogenica’s* putative starch utilization system loci consisting of a ten gene cluster. Gene locus tags are listed below each gene and known gene names are listed above each gene respectively.

To identify the specific polysaccharide utilization locus (PUL) responsible for starch degradation, we screened for PULs enriched in genes predicted to encode α-amylases and pullulanases. α-Amylases hydrolyze α-1,4 glycosidic linkages between glucose units in starch and its linear derivatives, such as maltotriose. In contrast, pullulanases can cleave α-1,6 glycosidic bonds found in branched starch structures and/or α-1,4 glycosidic bonds similar to α-amylases (9). The co-occurrence of these enzyme types within a locus is indicative of a PUL specialized for comprehensive starch degradation. Through this method, PUL3 was identified to have the most starch-implicated genes. PUL3 contains a predicted α-amylase encoded by gene RS00780, a pullulanase encoded by gene RS00785, and a second α-amylase encoded by gene RS00825. As a result, we conclude that PUL3 is the most likely candidate gene cluster responsible for the starch utilization system (sus) in *P. melaninogenica* (Fig 4).

### Candidate homologs were found for nine out of the ten genes in *P. melaninogenica’s* PUL3

Our objective was to determine the homologs of the putative starch utilization genes in *P. melaninogenica*. Our approach was to perform a BLASTp analysis using the amino acid sequences for each gene to obtain information on both the sequence and putative structural similarity. Due to the well characterized nature of *B. thetaiotaomicron’s sus* locus, we chose to first perform a BLASTp analysis on each PUL3 gene against each gene in *B. thetaiotaomicron’s sus* gene cluster (only the top hit is shown in Fig 5). If this did not yield a result, we then performed a BLASTp analysis against the entirety of the *B. thetaiotaomicron* genome in an untargeted search (Fig 5). If no homologs were found, we then looked in the closely related species *B. fragilis’ sus* gene cluster (only the top hit is shown in Fig 5). As with *B. thetaiotaomicron*, if no hits were found, we then performed an untargeted BLASTp search against the entire *B. fragilis* genome (Fig 5).

**Figure 5.**
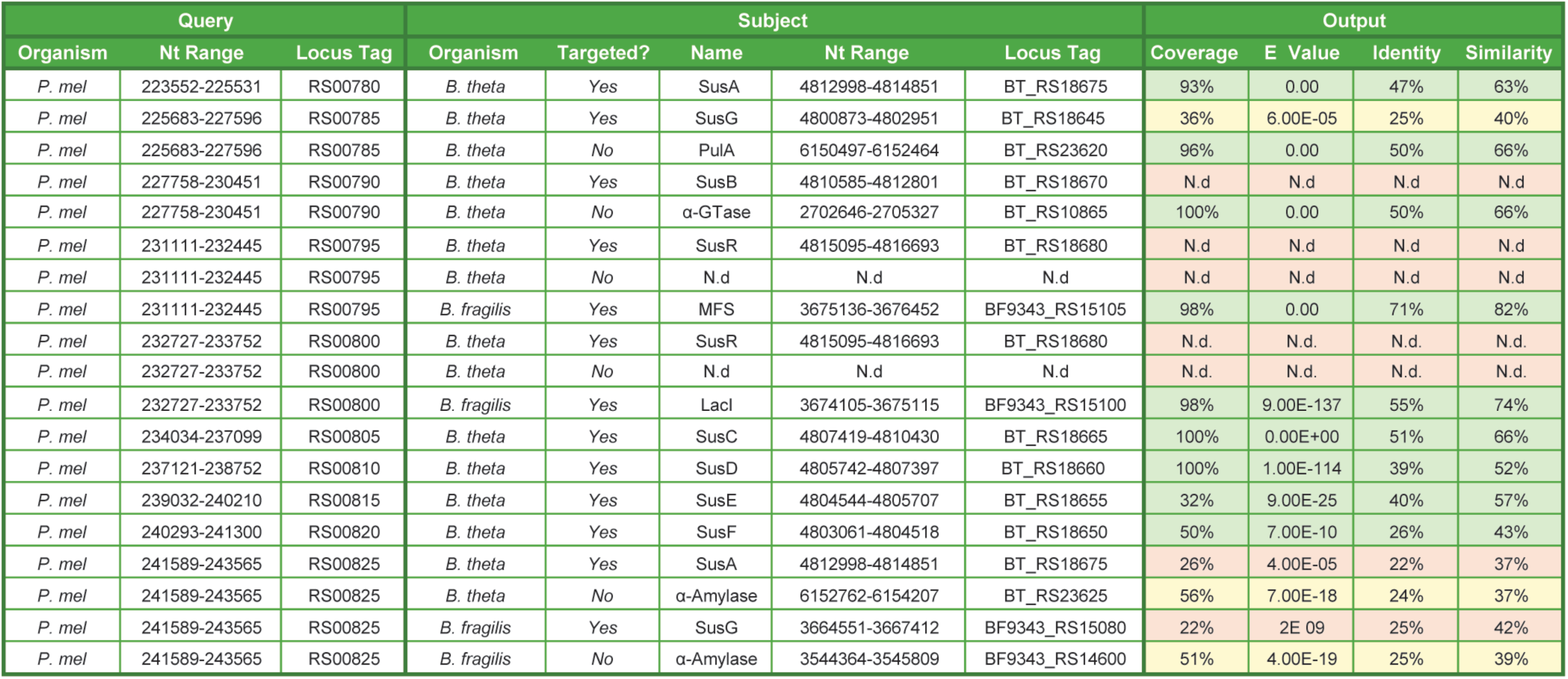
Candidate homologs of the putative sus locus genes in *P melaninogenica*. Table of amino acid BLAST results. The query column consists of *P melaninogenica’s* PUL 3 genes, starting with RS00780 and ending with RS00825. The subject column contains genes from *8. thetaiotaomicron* and *8. fragilis.* The Output column contains the BLAST results of each respective search. For each *P melaninogenica* gene, *we* first BLASTed against the best homolog candidate within the *8. thetaiotaomicron* sus gene cluster. This was categorized as a “targetted” search. If this did not yield a high coverage and similarity score, *we* then BLASTed the *P melaninogenica* gene against the entire *8. thetaiotaomicron* genome. This was categorized as a “untargetted” search. If this did not yield a hit with a high coverage and similarity score, the same process was carried out within *8. fragilis.* Candidate homologs with high sequence coverage and protein similarity are highlighted in green. If a hit had a relatively high coverage and similarity but a stronger homolog candidate was found, the hit is highlighted in yellow. BLAST alignments with no hits detected are listed as “n.d” and highlighted in red. The following reference sequenc­ es were used: *P melaninogenica* ATCC 25845 NC_014370.1; *8. thetaiotaomicron* VPI-5482 NC_004663.1; *8. fragilis* NCTC 9343 NC_003228.3.

Moving left to right across *P. melaninogenica’s* gene cluster, RS00780 was examined first. Within the *B. thetaiotaomicron* sus locus, a hit with 93% amino acid coverage and a protein similarity of 63% was found. The hit, named, is an α-amylase known to break down oligosaccharides in the periplasm. These results were supported by our finding that all of the known active site residues in *B. thetaiotaomicron’s* susA protein are predicted to be structurally conserved in RS00780’s amino acid sequence (Sup 1)(10). What’s more, the peptide sequence is predicted to encode a SPI signal peptide guiding translocation to the periplasm (Sup3). As a result, *susA* was determined to be a candidate homolog of gene RS00780 in *P. melaninogenica*.

We next examined gene RS00785 and found its amino acids sequence had a fair protein similarity of 40% to susG, but a coverage of only 36%. However, when a BLASTp analysis was carried out against the *B. thetaiotaomicron’s* genome, we found that protein PulA, a pullalanase, demonstrated a protein similarity of 66% and coverage of 96%. Therefore, homology to *PulA* was concluded.

Gene RS00790 did not have any hits detected within the *B. thetaiotaomicron* sus loci, notated as “N.d”. However, a 4-alpha glucotransferase (α-GTase), was found in the untargeted search with a coverage of 100% and protein similarity of 66%. Therefore, homology to the gene encoding this α-GTase was concluded.

Both RS00795 and RS00800 had no hits detected within both the *B. thetaiotaomicron* sus locus and *B. thetaiotaomicron* as a whole. However, both encoded proteins were found to have homologs in *B. fragilis’* sus locus gene cluster. The protein MFS had a 98% coverage and 82% protein similarity to RS00795’s amino acid sequence. The protein LacI had a 98% coverage and 74% protein similarity to RS00800. Therefore, gene homology was concluded.

Next looking at RS00805 and RS00810, both were found to have a high degree of coverage and similarity to *B. thetaiotaomicron’s susC* and *susD* respectively. We found that five of the six known active site residues in the encoded susD protein are structurally conserved in the RS00810 amino acid sequence (Sup 1)(11). Although the susC protein does not have active site residues for comparison, the predicted protein structure of RS00805 maintains the distinct transmembrane barrel morphology expected of susC (Sup 4). RS00805 and RS00810 are predicted to contain type I and type II signal peptides, respectively, and are directed to the outer membrane (Sup 3). Along with their domain architectures, this supports their annotation as *SusC* and *SusD* homologs.

When RS00815 and RS00820 amino acid sequences were aligned against lipoproteins susE and susF respectively, they had a fair degree of coverage and protein similarity (susE: 32% coverage, 57% similarity; susF: 50% coverage, 43% similarity). However, we found that the known susE and susF active site residues were both respectively conserved in RS00815 and RS00820, indicating a conserved function (Sup 1)(12). What’s more, the peptide sequences of both RS00815 and RS00820 are predicted to contain lipoprotein signal peptides and be secreted to the outer membrane (Sup 3). Finally, when examining conservation of the two known binding sites CBM Eb and CBM Ec in susE, we found that Eb was structurally conserved but Ec was not (Sup 1). However, Eb has been shown to be sufficient for alpha-glucan binding. RS00820’s peptide has structural conservation of 4 out of the 5 susF *a* binding site residues and 4 out of 5 susF *b* binding site residues (12). No residues in binding site *c* are present. However, similar to susE, susF does not need all three binding sites for glucan binding and therefore, we concluded homology of RS00815 and RS00820 to the *susE* and *susF* genes respectively.

Finally, RS00825 was examined for homology. The top hit within sus was to *susA*, however the amino acid coverage was only 26% and most of the active sites were not predicted to be conserved. When a BLASTp analysis was performed against all of *B. thetaiotaomicron*, the top hit was an α-amylase with 56% coverage and 37% protein similarity. The active sites are unknown for this α-amylase therefore an active site comparison was not possible. However, the predicted structure of this protein was aligned with that of RS00825 and three distinct domains were absent from the *B. thetaiotaomicron* α-amylase. This structural distinction can be seen in Figure 6b, where the unique domains are highlighted in blue. We found similar results when looking in *B. fragilis,* with a fair hit to an α-amylase but lacking three domains present in *P. melaninogenica’s* α-amylase. As a result, we concluded that the protein encoded by RS00825 is an α-amylase and that while it has some similarity to α-amylases within *B. thetaiotaomicron* and *B. fragilis,* no gene homolog was found.

**Figure 6.**
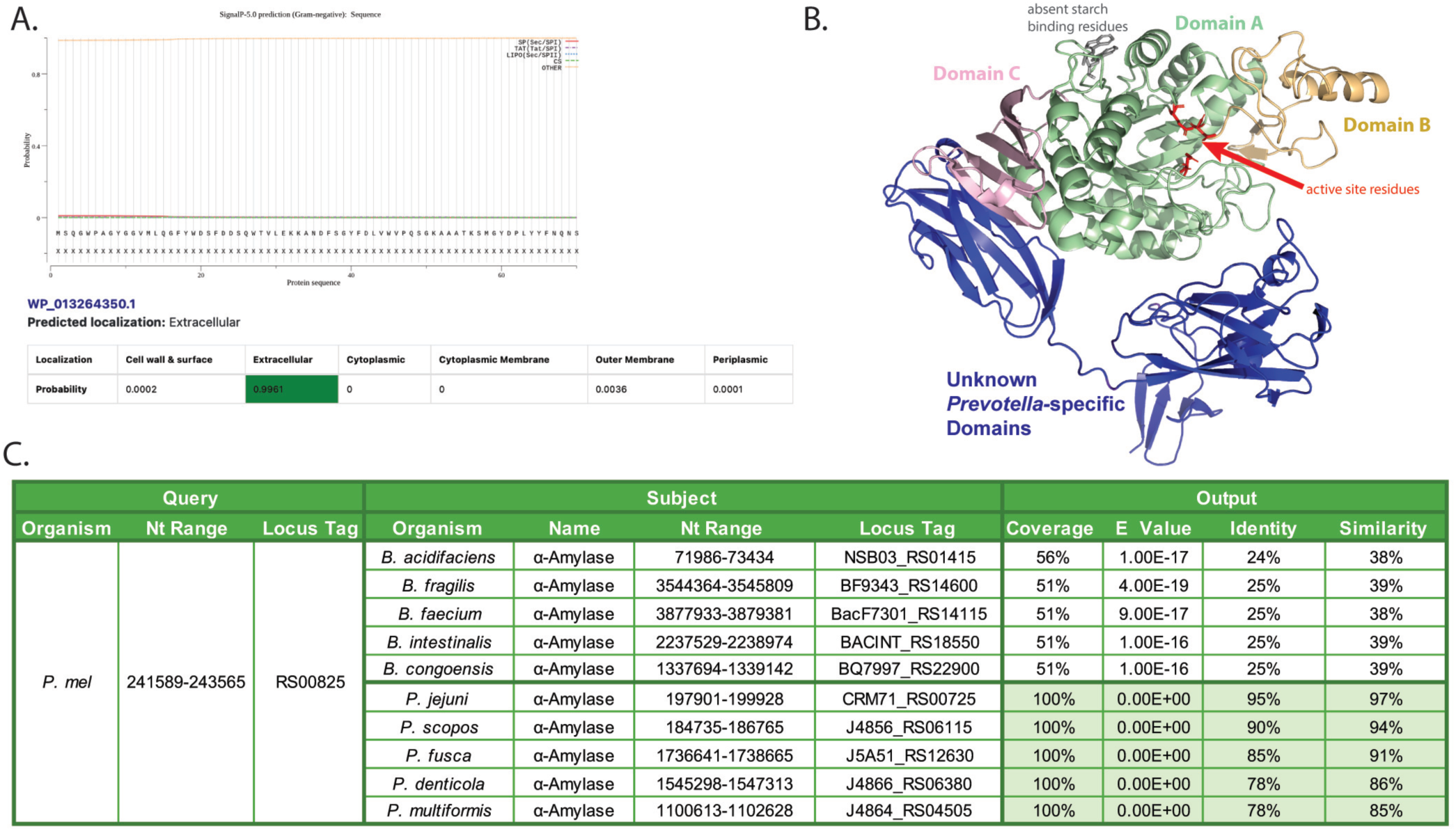
The RS00825-encoded a-amylase in *P melaninogenica* is a predicted extracellular protein with novel domains shared across the *Prevotella* genus and conserved active site residues. A. Predicted localization of the RS00825-encoded a-amylase. The top graph shows SignalP analysis, which predicts the probability of a signal peptide (SP) along the protein sequence (x-axis). No red, purple, or blue lines-indicative of SPs-are present, nor is a green line indicating a predicted SP cleavage site. The yellow “other” line represents amino acids not predicted to encode an SP. The bottom table presents DeepLocPro 1.0 predictions, which identify RS00825 as encoding a non-classical secretory protein with extracellular localization. B. Structural prediction and domain annotation of RS00825. The protein structure was modeled using AlphaFold and annotated based on the a-amylase of *Hordeum vulgare* (the most similar structure in the PDB). Domains A (green), B (yellow), and C (pink) are structurally conserved between *P melaninogenica* and *H. vulgare.* Domain A features a (l3a)a-barrel containing active site residues D179, E204, and D289 (labeled in red). Domain B is an irregular loop, and Domain C forms a five-stranded anti-parallel 13-sheet. Starch-binding sites present in *H. vulgare* (gray) are absent in *P melaninogenica.* Additional domains not found in *H. vulgare* or any *8acteroides* spp. are marked in dark blue. C. Sequence similarity analysis of the RS00825-encoded a-amylase with *Prevotella* and *8acteroides.* The BLAST query was *P melaninogenica* RS00825; subject sequences were from either *8acteroides* or *Prevotella.* The top five BLAST hits from each genus are listed, including gene name, nucleotide range, and locus tag. Output statistics are presented in the final column. Hits from Prevotella (highlighted in green) show significantly higher sequence similarity than those from *8acteroides.* Strains and reference genomes include: *8. acidifaciens* DSM 15896 (NZ_JAN­ JZR010000003.1), *8. fragi/is* NCTC 9343 (NC_003228.3), *8. faecium* CBA7301 (NZ_CP050831.1), *8. intestinalis* DSM 17393 (NZ_AB­JL02000008.1), *8. congoensis* Marseille-P3132T (NZ_FQXY01000011.1), *P jejuni* CD3:33 (NZ_CP023863.1), *P scopos* W2052 (NZ_CP072390.1), *P fusca* W1435 (NZ_CP072370.1), *P dentico/a* F0115 (NZ_CP072374.1), and *P multiformis* F0096 (NZ_CP072357.1).

All together, we were able to identify homologs for nine out of the ten genes present in *P. melaninogenica’s* starch utilization loci. Notably, some of the homologs were found outside of the *B. thetaiotaomicron* or *B. fragilis* sus loci.

### Gene RS00825 encodes the putative extracellular α-amylase which features additional domains that are conserved within the *Prevotella* genus but absent in *Bacteroides*

Our objective was to predict the localization and structure of the α-amylase encoded by gene RS00825 (hereby called *AmyA*). These findings will inform the enzyme’s putative function and its role in starch degradation. To assess whether RS00825 encodes a secreted protein, we first used SignalP to analyze the amino acid sequence for canonical signal peptides, including secretory, lipoprotein, and TAT signals. No such signal peptide was predicted (Fig 6a). To evaluate the potential for non-classical secretion, we then applied DeepLocPro 1.0, a neural network-based tool. DeepLocPro predicted AmyA to be extracellular with a probability of 0.99 out of 1, based on its amino acid sequence (Fig 6a). Therefore, we concluded that AmyA is an extracellular protein excreted through a non-canonical mechanism.

To investigate the structural features of *P. melaninogenica*’s AmyA, its structure was first predicted using AlphaFold 3 and then aligned to the crystal structure of *Hordeum vulgare* α-amylase, the closest homolog available in the PDB. This comparative analysis revealed conservation of several key structural domains (Fig 6b). Domain A contains a (βα)₈ -barrel and includes the catalytic residues D179, E204, and D289. Domain B forms an irregular loop that coordinates three Ca²⁺ binding sites, while Domain C adopts a five-stranded anti-parallel β-sheet. Notably, the starch-binding residues W276 and W277 found in Domain A of *H. vulgare* are not conserved in *P. melaninogenica*. However, AmyA contains additional β-sheet domains that are absent in *H. vulgare*. A search of the amino acid sequence corresponding to these regions in the PDB yielded no structural hits.

To assess the conservation of this unique α-amylase structure, we performed BLASTp searches of AmyA’s amino acid sequence against the *Prevotella* and *Bacteroides* genera respectively (Fig 6c). The results revealed high sequence conservation within *Prevotella*, with the top five hits exhibiting an average protein similarity of 90.6% and 100% coverage. In contrast, the top five *Bacteroides* hits exhibited an average protein similarity of 38.6% and 52% coverage, indicating limited conservation in this closely related genus.

### Gene RS00875 is a putative extracellular and periplasmic protein

Our objective was to characterize the localization and structural features of the RS00875-encoded pullulanase we are calling PulA. As with AmyA, these analyses aim to inform its putative function and potential role in starch degradation. PulA was predicted by the PULdb to be a member of the glycoside hydrolase family 13, subfamily 14. As a GH13_14 protein is not known to be present in any *Bacteroides* starch utilization loci, the presence and potential role of PulA within the *P. melaninogenica* Sus system represents a novel finding. To begin assessing its function, we first examined the protein sequence for the presence of a signal peptide, which could provide insight into its localization. Utilizing SignalP-5.0, a secretory signal peptide type I was predicted to be present between amino acids 1-21 of the protein (Fig 7a). Furthermore, a cysteine was present at the SP cleavage site (Fig 7a). These results suggest that the protein is secreted via a canonical secretory signal peptide. To predict the subcellular localization of PulA, we used DeepLocPro 1.0 and found that PulA is predicted to be localized to both the extracellular matrix and the periplasm, with a probability of almost 50% respectively (Fig 7a and Sup3). We therefore concluded that PulA contains a canonical secretory signal peptide which allows for its localization to the periplasm and extracellular space.

**Figure 7.**
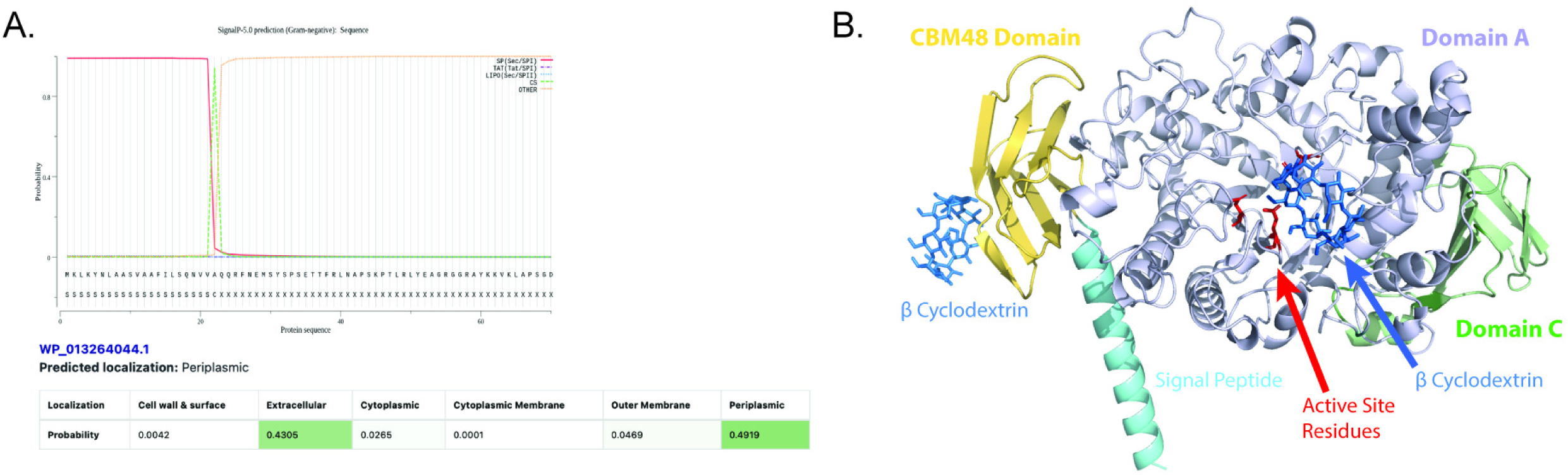
The RS00785-encoded pullulanase in *P. melaninogenica* is a predicted extracellular and periplasmic protein with conserved active site residues. A. The presence of a signal peptide encoded by *P melaninogenica’s* RS00785-encoded PulA with SignalP 5.0. The predicted cysteinated secretory signal peptide I (SPI) encoded by the 5’ 21 amino acid sequence is depicted as a red line. The signal peptide cut site is depicted as a green dotted line with a spike between amino acids 21-23, indicating the local where the signal peptide is predicted to be cleaved. The yellow line, labeled *other* demonstrates the remaining amino acids of the protein. The C residue adjacent to the cut site is characteristic of a peptide sequence post-cleavage. B. The structure of *P melaninogeni­ ca’s* RS00785-encoded PulA was predicted with Alphafold and annotated based on the PulA of *Paenibacil/us barengoltzii* (the most similar structure in the PDB with substrate binding). Domains A, C and CBM48 were conserved, with active site residues D320, E349 and D449 labeled in red. Co-crystalized cyclodextrin bound to the starch binding sites in CBM48 and the active site are depicted in blue. The encoded signal peptide pre-cleavage is depicted in light blue.

**Figure 8.**
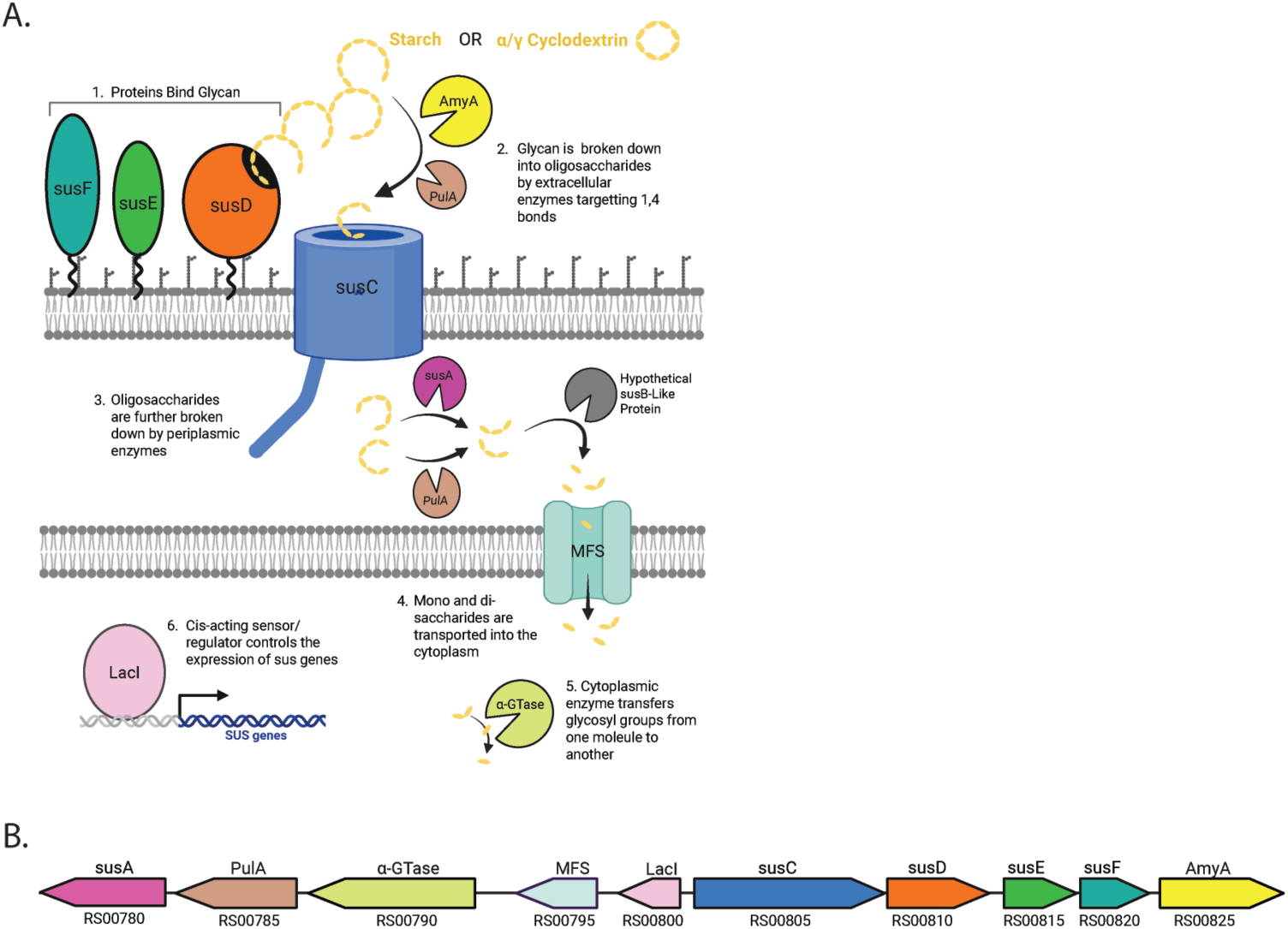
The putative starch utilization system in P. melaninogenica. A. Model of the predicted starch utilization system in *P. melaninogenica*. 1. Starch or starch derivatives such as cyclodextrin bind to the proteins susF, susE and susD. 2. PulA and/or a-amylase AmyA perform the first breakdown step, cleaving the glycan into olygosaccharides which are transported into the periplasm by susC. 3. Oligosaccharides are further broken down into maltotriose by either susA and pulA before subsequent breakdown by a hypothetical susB. 4. Glucose and maltose are transported into the cytoplasm by the MFS protein. 5. 4-alpha-glucotransferase breaks the a-1,4 bonds of maltose ans transfers a glucosyl groupt to glucose or maltose. 6. The transcription factor Laci regulates the expression of sus genes. B. P. *meianinogenica*’s putative sus gene cluster with genes labeled according to their predicted function. Homologs of *B. thetaiotaomicron* or *B. fragilis* sus genes are depicted in the same color.

We next aimed to examine the structure of PulA. Like with AmyA, our approach was to identify the protein crystal structure in the PDB database with the highest similarity to PulA and use this as a structural reference. Our top hit was a Pullulanase in *Paenibacillus barengoltzii* which had a sequence coverage of 89% and protein similarity of 61% (13). Notably, the Pullulanase was co-crystalized with cyclodextrin, enabling an additional analysis of the potential starch binding sites of *P. melaninogenica’s* PulA protein. The Pullulanase protein contains three domains: The catalytic site, the CBM48 Domain and Domain C. The catalytic site of *P. barengoltzii* can bind both cyclodextrin and longer oligosaccharides and is defined by the presence of residues D350, E379 and D470 (13). All three active site residues were found to be conserved in *P. melaninogenica’s* PulA, supporting PulA’s ability to hydrolyze starch bonds at this site. PulA’s active site residues and the superimposed cyclodextrin substrate binding can be seen in Figure 7b. The CBM48 domain of *P. barengoltzii’s* pullulanase can only bind cyclodextrin and is defined by F63 and W111. *P. melaninogenica’s* PulA does not contain these residues, however, the two cyclodextrin binding residues involved in sugar-aromatic stacking interactions are known to vary between organisms. The superimposed cyclodextrin binding at the CBM48 Domain of *P. melaninogenica’s* PulA can be seen in Figure 7b. Domain C was found to have a novel cyclodextrin binding ability in *P. barengoltzii’s* pullulanase which is dependent on the presence of a short FGGEH motif. However, similar to other pullulanases like KPP and HvLD(14,15), *P. melaninogenica’s* PulA contains long motifs in the 5 and 6 Beta-strands which prevent sugar binding (Fig 7b). Based on these results, we conclude that *P. melaninogenica’s* PulA, encoded by gene RS00875, likely contains the residues necessary for cyclodextrin binding and breakdown.

## DISCUSSION

We developed and validated a basal medium for carbon-utilization assays in *P. melaninogenica* by assessing growth on key central carbohydrates. The development of a growth medium that enables significant growth differences in response to carbon supplementation overcomes a key methodological barrier and establishes a platform for future functional studies. Using this medium, we mapped *P. melaninogenica*’s putative starch utilization system and identified three proteins—AmyA, PulA, and aGTase—that are absent from any *Bacteroides* sus model. This divergent architecture indicates lineage specific starch utilization pathways and provides a novel framework for studying *Prevotella* carbohydrate metabolism. Collectively, these findings provide valuable insights into how this genus may adapt to nutrient variability across diverse host-associated environments.

When testing the carbon utilization capacity of *P. melaninogenica*, we found it efficiently utilized glucose, fructose, lactose, sucrose, maltose, maltotriose, and cyclodextrin—a starch mimic. Given the limited knowledge in the literature regarding small sugar uptake mechanisms in *Prevotella* and the closely related genus *Bacteroides*, we turned to the extensively studied model organism *Escherichia coli* for comparative insights. In *E. coli, s*mall molecular weight sugars like glucose are transported non-specifically through the outer membrane by porins. Once in the periplasm, the sugars are transported through the inner membrane by several systems, including phosphoenolpyruvate-dependent phosphotransferase system (PTS), the ATP-dependent cassette (ABC) transporters, and the major facilitator superfamily proton symporters (MFS) (16,17). Whether this mechanism is conserved in *P. melaninogenica* remains unknown, highlighting the need for further investigation. In contrast, the utilization of starch has been extensively studied in the *Bacteroides* genus, with *B. thetaiotaomicron* representing the first fully characterized Sus system. Given our findings that *P. melaninogenica* can utilize cyclodextrin, a starch mimic, investigating its starch utilization system (sus) represented a logical next step.

Within *Bacteroides*, many polysaccharides have their own utilization system, each encoded by a polysaccharide utilization loci (PUL) gene cluster (18–20). The starch utilization system (sus) in *B. thetaiotaomicron* is encoded by PUL66, an eight gene cluster called the sus loci encoding susRABCDEFG. susDEF are members of the outer membrane which recognize and bind to starch, sequestering it at the bacterial membrane. The outer membrane bound α-amylase susG then degrades starch into smaller maltooligosaccharides by targeting its alpha 1,4 bonds (21,22). These starch derivatives are then transported into the periplasm by susC, a tonB dependent β-barrel porin(23,24). Maltooligosaccharides are then further broken down by the α-amylase susA and a-glucosidase susB, resulting in single glucose units. Glucose is then transported into the cytoplasm. The sus loci expression is regulated by the transmembrane protein susR which senses maltose in the periplasm and upregulates gene expression (21).

Our investigation of *P. melaninogenica*’s starch utilization system revealed a distinct gene cluster containing several components not found in the well-characterized starch utilization system in *B. thetaiotaomicron* and three encoded proteins which are not found in any *Bacteroides* sus loci. The cluster shares some conserved elements, such as proteins susC, susD, susA, susE, and susF. However, the cluster also encodes MFS and LacI which are absent in *B. thetaiotaomicron*, lacks *B. thetaiotaomicron’s susR* and contains three novel sus proteins—amyA, pulA, and aGTase—not thought to be present in any *Bacteroides* starch utilization system. These unique features point to a divergent system architecture that likely reflects lineage-specific adaptations for carbohydrate acquisition. Collectively, these findings broaden our understanding of *P. melaninogenica*’s metabolic strategies and suggest mechanisms by which it may be specially equipped to thrive in the nutrient-variable conditions of mucosal environments, such as the human respiratory tract.

Gene RS00825 is predicted to encode the α-amylase AmyA, a protein absent from the *Bacteroides* genus but broadly conserved across *Prevotella*, suggesting divergence in the mechanism of starch utilization. We found that AmyA is predicted to be found in the extracellular space, consistent with its role in hydrolyzing α-1,4 glycosidic bonds in starch and related oligosaccharides. In *B. thetaiotaomicron*, the α-amylase enzyme SusG is anchored to the outer membrane alongside the SusCDEF complex, facilitating coordinated starch degradation at the cell surface. While AmyA may serve a similar function in *P. melaninogenica*, its dissimilar structure and lack of a predicted membrane anchoring suggest a varied rate of catalysis. This feature may enable AmyA to act independently of membrane sequestration, potentially increasing the efficiency of starch degradation and representing a distinct, possibly more flexible mechanism of extracellular carbohydrate processing compared to members of the *Bacteroides* genus.

The gene RS00790 is predicted to encode a 4-α-glucotransferase (αGTase) that, while present in *Bacteroides*, is not associated with any characterized polysaccharide utilization loci (PULs), including those involved in starch metabolism (25). In contrast, this enzyme is predicted to be broadly distributed across *Prevotella* PULs. Notably, it is found exclusively within PULs that also contain high densities of amylases and pullulanases, suggesting that its presence may be specific to the starch utilization loci in *Prevotella* species. These patterns indicate that the αGTase may represent a conserved and defining component of the starch utilization system in this genus. Although the identified *B. thetaiotaomicron* αGTase (BtαGTase) homolog is not part of the sus locus, its characterized function may offer insights into the potential role of αGTase activity in *P. melaninogenica*’s starch metabolism. BtαGTase catalyzes disproportionation reactions through the repeated hydrolysis and resynthesis of α-1,4-glycosidic linkages (26). The enzyme may use starch and amylopectin as donor substrates and may use a variety of maltooligosaccharides—including maltose, maltotriose, maltotetraose, maltopentaose, maltohexaose, and maltoheptaose—as acceptors. These reactions can occur both intermolecularly and intramolecularly. In the latter case, intramolecular transglycosylation can lead to the formation of cyclic glucans known as cycloamyloses, which are structurally similar to cyclodextrins (26). The role of this enzyme could represent an important addition to current models of starch metabolism. If starch is used as the donor substrate for transglycosylation, the *sus* paradigm suggests that the enzyme should be localized to the outer membrane. Surprisingly, when the amino acid sequence of both BtαGTase and PmαGTase was analyzed, neither were predicted to be secreted. When further analyzed with DeepLocPro 1.0, the protein was again predicted to be cytoplasmic. While the transglycosylation activity of αGTase likely contributes to starch metabolism in *P. melaninogenica*, its specific role within the cytoplasm remains unresolved. The conservation of αGTase across *Prevotella* species, coupled with its absence from the *sus* loci of *Bacteroides*, suggests that it may represent a lineage-specific adaptation to the ecological pressures associated with diverse host environments.

The third gene characteristic of a *P. melaninogenica* starch utilization system is RS00785. This gene is predicted to encode the Pullulanase PulA. While a similar pullulanase is predicted to be present in some *Bacteroides* PULs, the PULs lack the characteristics of a sus loci. By contrast, PulA in *Prevotella* is predicted to be present exclusively in PULs characteristic of sus loci, suggesting that PulA may be a defining feature of the *Prevotella* starch utilization system. Interestingly, PulA is predicted to undergo SPI-type excretion to both the extracellular and periplasmic space. Due to PulA’s predicted structural conservation, it is likely that PulA retains the ability to bind starch at the CBM20 domain and cleave glycosidic bonds in the active site. However, due to the results of Figure 2, it is unlikely that *P. melaninogenica* can hydrolyze the α 1,6 bond, a characteristic function of pullulanases. Notably, it has been found that some pullulanases present in Bacteroides spp. like *B. thetaiotaomicron* contain pullulanases which demonstrate α 1,4 hydrolysis activity. This activity in PulA would make sense in the context of starch degradation in the extracellular and periplasmic space of the *P. melaninogenica*.

Like with AmyA, PulA is not predicted to be anchored to the outer membrane, unlike proteins in the canonical sus pathway. Whereas in *B. thetaiotaomicron* the α-amylase SusG is the only starch degrading enzyme on the cell surface, we proposed that in *P. melaninogenica*, both α-amylase AmyA and pullulanase PulA are present and enable rapid α 1,4 bond degradation. This dual approach could enable a more flexible and efficient degradation of the starch substrates, demonstrating a unique adaptation for metabolic optimization within *P. melaninogenica.* The presence of PulA in the periplasm would support this hypothesis. With a pullulanase in the periplasm, oligosaccharides produced by AmyA and PulA α 1,4 degradation would be further broken down by α-amylase susA and PulA in the periplasm. The presence of both an α-amylase and pullanase in both the extracellular and periplasmic space would act as a more robust two-stop degradation which may result in a more rapid sugar intake.

Compared to smaller molecular weight carbon sources, *P. melaninogenica* exhibits a slower replication rate when grown on cyclodextrin. While basal expression of the *sus* system is likely maintained, the full upregulation of sus genes in response to complex carbohydrates such as starch or cyclodextrin may require additional time, contributing to the observed growth lag. In contrast, smaller sugars such as maltose or maltotriose may be imported via *sus*-independent pathways, potentially through passive diffusion across outer membrane porins, allowing for more immediate utilization. We hypothesize that while outer membrane uptake may proceed independently of *sus*, the transport of small molecules across the inner membrane may still be under *sus*-dependent MFS and LacI regulatory control, irrespective of starch availability. Supporting this model, the genomic organization of the *sus* cluster shows a bidirectional arrangement: approximately half of the genes, including those encoding the MFS transporter and LacI, are transcribed in one direction, while the remainder are transcribed in the opposite. This configuration suggests the presence of two distinct operons, which could allow for differential gene regulation. Such a mechanism may permit constitutive or semi-constitutive expression of MFS and LacI, while other sus genes are induced only in response to complex polysaccharides. This mechanism of action would allow for a flexibility in metabolic response reflective of P. melaninogenica’s varied niche environments.

The ecological significance of P. *melaninogenica*’s distinctive starch utilization pathways is particularly relevant given its ability to colonize both the oral cavity and the respiratory tract. These two environments present markedly different nutritional landscapes. In the mouth, dietary intake provides a continual influx of carbohydrates, proteins, and fats, in addition to host-derived mucins. In contrast, the respiratory tract offers a more limited and competitive nutrient environment, where available carbon sources are primarily restricted to host-derived compounds such as mucins and other substrates transported through bidirectional airflow. To successfully inhabit both niches, *P. melaninogenica* must possess a highly adaptable metabolic profile capable of modulating carbon source utilization in response to environmental variability. These novel components within the *sus* system likely reflect evolutionary adaptation to the host niche, supporting the idea that host association has shaped metabolic specialization.

This study provides a foundational framework for the future molecular validation of *sus* system components in *P. melaninogenica*. Through preliminary testing of central carbon sources, identification of putative starch-utilization loci, and computational predictions of gene function, this work offers critical insights into the organism’s metabolic potential. Although the absence of a genetically tractable *P. melaninogenica* strain currently limits direct functional validation, developing such a system remains a key priority for advancing experimental studies. In the interim, future directions include qPCR-based analysis to assess gene expression in response to starch supplementation, which may reveal transcriptional regulation patterns. Additional investigation into co-regulated *sus* genes could help define operon structures and distinguish responses to maltooligosaccharides versus starch. Furthermore, the development of a defined basal medium for carbon utilization assays contributes to a valuable toolkit for functional and ecological studies. Collectively, these efforts establish a robust platform for elucidating starch metabolism in *P. melaninogenica* and understanding its adaptation to host-associated environments.

## METHODS

### *P. melaninogenica* reconstitution and culture conditions

*Prevotella melaninogenica* 25845 was purchased from ATCC as a frozen culture. Bacteria was grown up in Anaerobe systems BHI broth under anaerobic conditions. Gram stain and 16S sequencing was performed to confirm ID.

### Basal media development

A basal medium was formulated to include essential trace elements sufficient to support the optimal growth of *P. melaninogenica*. To enable controlled carbon utilization studies, the medium was designed with minimal levels of carbohydrates and amino acids, thereby making carbon availability the limiting factor for growth. As a result, the medium supports only low-level growth in the absence of supplementation. However, when supplemented with an additional carbon source that *P. melaninogenica* can metabolize, the medium permits a marked increase in growth, enabling assessment of substrate-specific utilization.

The low-level carbon sources in BBM medium are provided through the inclusion of brain heart infusion broth, proteose peptone, and yeast extract. Sodium chloride and sodium phosphate dibasic are added to maintain optimal osmotic pressure and pH balance. To support the growth of anaerobic organisms, L-cysteine hydrochloride is included as a reducing agent, and resazurin is added as a redox indicator. Hemin and vitamin K₁ , both previously reported as essential for *P. melaninogenica* growth, are also incorporated (Table 1). After all components are combined, the volume is adjusted to 1 liter with water and thoroughly mixed. The final pH should range between 7.0 and 7.5; pH adjustment with NaOH is not recommended, as it may interact adversely with an unidentified component of the medium. Because the medium is sensitive to heat and light, it should be filter-sterilized and stored protected from light. Prior to use with anaerobic cultures, allow the medium to reduce under anaerobic conditions for 48 hours.

### Assaying Carbon Utilization

Carbon utilization assays were performed using the BIOLOG^©^ Phenotype MicroArray carbon panel. The carbon panel consists of two pre-configured 96 well plates named PM1 and PM2 which contain a different carbon source lyophilized within each respective well, enabling the simultaneous assessment of 190 distinct carbon substrates. Well A1 on each plate is left empty to serve as a negative control. *P. melaninogenica* was grown overnight in its preferred rich medium (Anaerobe Systems BHI broth©), then harvested by centrifugation and resuspended in BBM. To achieve a final inoculum concentration of 1.66% and a starting optical density (OD) of 0.05 per well, the bacterial culture was adjusted to an OD of 3.00. For each well, 2.33 µL of the culture was mixed with 137.67 µL of BBM, resulting in a total volume of 140 µL. Plates were sealed with SealPlate (Excel Scientific©), and friction was applied to ensure a tight, airtight seal—an essential step for maintaining anaerobic conditions after removal from the anaerobic chamber. Sealed plates were removed from the anaerobic chamber and loaded into the ODIN© plate reader, where OD was measured every 10 minutes for 24 hours. Although readings were taken at 600 and 740 nm, only 740 nm data were analyzed to avoid interference from carbon source absorbance and more accurately reflect cell density.

### Genome annotation and glycolytic pathway analysis

To investigate the presence of glycolytic components in *P. melaninogenica*, we first referenced the KEGG pathway database to identify genes associated with the canonical glycolysis pathway. We then annotated the genome of *P. melaninogenica* strain ATCC 25845 using the DRAM (Distilled and Refined Annotation of Metabolism) pipeline. DRAM integrates multiple databases—including KEGG, UniRef90, and MEROPS—and employs MMseq2 to predict open reading frames, categorizing genes into functional groups such as energy metabolism, transporters, carbon utilization, organic nitrogen, and miscellaneous functions. Genes from the KEGG reference glycolysis pathway were cross-referenced with the DRAM output to assess their presence or absence in the genome of strain ATCC 25845. This comparative analysis confirmed the presence of core glycolytic genes.

### Polysaccharide utilization locus prediction

The PUL database was used to identify candidate PULs encoding the starch utilization system of *P. melaninogenica*.

### Protein Homolog Predictions

Protein similarity was first assessed using BLASTp to determine percent coverage, E value, amino acid identity and protein similarity. ClustalOmega Multiple Sequence Alignment was used to visualize amino acid alignments and the ClustalW with character output was chosen to highlight conserved structural features. All protein structures were predicted using AlphaFold3 and analyzed on PyMol.

### Protein Localization Predictions

Protein localization was predicted using SignalP 5.0 and DeepLocPro 1.0. SignalP 5.0 predicts the presence of N-terminal SPI, Lipoprotein or TAT signal peptides in gram negative bacteria. SignalP 5.0 is based on a deep convolutional and recurrent neural network architecture including a conditional random field. DeepLocPro 1.0 predicts the presence and subcellular localization of non-canonical secretory proteins. However, it is also effective in predicting the localization of canonical secretory proteins. DeepLocPro 1.0 is a a neural network-based tool trained on UniProt and ePSORTdb proteins with experimentally validated subcellular localizations.

## CONCLUSIONS AND OUTLOOK

*P. melaninogenica* is a highly abundant bacteria of the human respiratory tract that remains poorly understood compared to other respiratory microbes. Colonization within the nutrient rich environment of the mouth and less habitable environment of the lower respiratory tract indicates a sophisticated metabolic system equipped to deal with large fluctuations in nutrient availability. In this study we present a basal media recipe which enables carbon utilization testing of *P. melaninogenica*. Applying this tool, we demonstrate the utilization of dietary carbohydrates with α 1,4, β 1,4 and α 1,2 glycosidic bonds, like the starch mimics α and γ-cyclodextrin. Going a step further, we then identified the putative starch utilization locus and the AmyA, PulA and GTase components absent from the *Bacteroides* genus. This work significantly advances our understanding of the metabolic capabilities of *P. melaninogenica*. By beginning to uncover its carbohydrate utilization and starch breakdown system, we’ve paved the way for deeper insights into microbial colonization within the respiratory tract.

## ACKNOWLEDGEMENTS

C.E.A. was supported by the following NIH Training Grant: Michigan Predoctoral Training in Genetics (T32GM149391). K.S. was supported by The University of Michigan Rackham Merit Fellowship. This work was also supported by the University of Michigan. We thank our colleagues Dr. Nicole Koropatkin and Dr. Eric Martens for providing their expertise and guidance.

